# Pathogenic *MAPK8IP3* variants drive distinct motor and behavioral phenotypes in humans and mice

**DOI:** 10.64898/2026.07.20.739682

**Authors:** Camerron M. Crowder, Lyndsay R. Watkins, Alexa Geltzeiler, Priya Patel, Judit Perez-Ortiz, Danielle K. Schmidt, Christopher Schmitz, Jalen Aguilar, Shraddha Ghanta, Wendy. K. Chung, Swetha Gowrishankar, Laura Lambert

## Abstract

Pathogenic variants in *MAPK8IP3* (JIP3) cause Neurodevelopmental Disorder with or without variable Brain Abnormalities (NEDBA), characterized by cognitive impairment, global developmental delay, motor dysfunction, abnormal muscle tone, behavioral dysregulation including autism and attention deficit hyperactivity disorder (ADHD), seizures, and structural brain abnormalities. While more than 30 largely *de novo MAPK8IP3* variants have been reported, the functional impact of variants across JIP3 protein structural domains is poorly defined. To address this knowledge gap, we compare clinical features of individuals with a truncating (p.E27X) or one of two missense (p.R578C and p.R1146C) variants from distinct JIP3 functional protein domains to corresponding knock-in mouse models. Our findings showed that all individuals, regardless of variant type, exhibited delays in language and gross motor function, but individual variants were associated with distinct motor, cognitive, and psychiatric symptoms. Corresponding homozygous p.E27X and p.R1147C variant mice resulted in embryonic lethality, consistent with essential roles for JIP3 in early neurodevelopment. Behavioral characterization of viable heterozygous mice revealed variant-specific locomotor, motor coordination, and hindlimb clasping defects that closely recapitulate clinical observations. Heterozygous p.R579C mice exhibited reduced locomotion, impaired motor performance, and hypertonia-like clasping, mirroring human motor deficits. In contrast, p.R1147C mice displayed hyperactivity, hindlimb clasping, and decreased brain weight, paralleling human clinical features. Together, our findings demonstrate that while distinct *MAPK8IP3* variants lead to some shared phenotypes, they are also associated with distinct phenotypes that could reveal domain-specific aspects of JIP3 function. This work establishes the first domain-resolved *in vivo* rodent models of NEDBA and provides a validated translational platform for mechanistic investigation and preclinical therapeutic testing.

## INTRODUCTION

*MAPK8IP3* (*Mitogen-Activated Protein Kinase 8 Interacting Protein 3*), encodes JNK Interacting protein 3 (JIP3), which functions as both a scaffold for the c-Jun NH_2_-terminal kinase (JNK) signaling pathway and as an adaptor linking organelle and vesicular cargoes to the motor proteins kinesin and dynein. Through these roles, JIP3 regulates bidirectional, microtubule-based cargo transport in neurons (1–6). Orthologs of JIP3 have been described across model organisms under species-specific names (JSAP1, UNC-16, Syd), and extensive work in these systems has established JIP3 as an evolutionarily conserved regulator of axonal transport (7).

Specifically, JIP3 is required for dynein-mediated retrograde trafficking and modulates kinesin-driven anterograde transport, ensuring proper positioning and long-range movement of endosomes, autophagic-and lysosomal intermediates, lysosomes, and signaling vesicles along axons (3,8–16). In mice, homozygous JIP3 knockouts exhibit severe telencephalic commissure defects, respiratory failure, and perinatal lethality (2), while heterozygous JIP3 mice are fully viable without appreciable defects. Cellular studies further demonstrate that JIP3 loss results in pathological accumulation within axonal swellings, impairing axonal autophagic-lysosomal clearance (11,13,15,16). Loss of JIP3 is linked to amyloidogenic APP processing and its haploinsufficiency enhanced amyloid plaque pathology in a mouse model of Alzheimer’s disease, linking loss of JIP3 function to neurodegenerative diseases (3,11,12,16). Beyond its established role in organelle transport in mature neurons, JIP3 function has been linked to aspects of neurodevelopment, including early axonal patterning, polarity establishment, and neurite morphogenesis (3,8,12,17).

Pathogenic and likely pathogenic variants in *MAPK8IP3* (JIP3) cause a spectrum of neurodevelopmental disorders collectively known as Neurodevelopment Disorder with or without variable Brain Abnormalities (NEDBA) (18). Affected individuals have cognitive impairment, global developmental delay, motor and balance difficulties, hypo or hypertonia, autism-spectrum disorder (ASD), attention-deficit/hyperactivity disorder (ADHD), seizures, microcephaly and variable structural brain abnormalities (18–20). To date, 32 mostly *de novo*, heterozygous pathogenic and likely pathogenic *MAPK8IP3* variants have been reported that localize across four major functional domains of the JIP3 protein (17). These include predominantly missense variants (62.5%) and predicted loss-of-function variants (34.4%) including nonsense, frameshift, and deletion variants (18). Missense variants may result in toxic gain of function, as reported in human-derived patient cells harboring the p.R578C variant (21) and generally manifest with more severe phenotypes (17). Haploinsufficiency variants are associated with milder phenotypes (17). Development of animal models that replicate patients symptoms is a critical first step to defining the pathophysiological mechanisms of JIP3 dysregulation, determining variant-specific functional, physiological consequences, and enabling preclinical testing of targeted interventions. These distinct mechanisms have important implications for therapeutic development, as toxic-gain of function variants should be targeted by knockdown strategies, such as antisense oligonucleotides (ASOs), whereas loss-of-function variants will likely require approaches that restore or augment JIP3 protein expression (18,21).

In this study, we compare neurological symptoms in humans across three *MAPK8IP3* variants spanning distinct JIP3 functional domains using clinician-administered motor and cognitive assessments (17). To further investigate the domain-specific functional consequences of these JIP3 variants, we obtained corresponding knock-in mouse models (E27X, R579C, and R1147C) and systematically characterized behavioral and motor phenotypes relevant to the neurological symptoms observed in humans. Collectively, these results clarify conserved genotype-phenotype relationships and establish robust, quantifiable behavioral metrics suitable for future preclinical testing of therapeutic strategies. Characterization of these models also provides a foundation for future molecular, cellular, and electrophysiological studies examining how distinct JIP3 variants affect the development and function of specific brain regions and neuronal populations.

## METHODS

### Patient neurological data

Clinical assessments were conducted in-person, as previously described (18). Clinical parameters were evaluated in a 7-year old male individual with a truncating p.E27X (E27X) variant in the RH1 domain, five individuals (2 males and 3 females age 7-24 years)with the missense p.578C (R578C) variant in the RH2 domain, and twelve individuals (7 males and 5 females age 3-27 years) with the missense p.R1146C (R1146C) variant in the WD40 domain. Motor function was assessed by a physical therapist using the Gross Motor Function Classification System with the Gross Motor Function Measure-88 converted to GMFM-66 scores (22,23). Upper limb function was measured using the Box and Block Test and nine-hole peg test (24,25). Gait was measured using the 10-meter walk run test, 6-minute walk test, and quantitatively measured on a Zeno instrumented walkway (26–28). Cognitive assessments utilized one of the following, dependent on age and ability: the Bayley Scales of Infant and Toddler Development (4^th^ edition), the Differential Ability Scales (2^nd^ edition), the Wechsler Abbreviated Scale of Intelligence (2^nd^ edition) (29–32).

The canonical human and mouse *MAPK8IP3* transcripts are listed in **Table 2**. Protein alignments between human and mice with both 1337 amino acid sequences were performed using the multiple sequence alignment tool in CLUSTALW (v2.1) (**Supplemental Figure 1**). Genotyping of mouse lines show nucleotide changes respective of stop codon (c.79 G>T, p.Y37X) and missense variant changes (c.1735 C>T, p.R578C and c.3439 C>T, p.R1146C) and restriction enzyme silent nucleotide changes required for genotyping (**Supplemental Figure 2**).

### Animal husbandry

Mice with variants engineered into the murine *MAPK8IP3* locus (p.E27X, p.R579C, p.R1147C (ENSMUST00000088345.12), corresponding to human patient variants (NM_015133) on the FVB background strain) were obtained from Charles River where they were bred and genotyped. Mice were originally generated by the Dr. Matthew Anderson Laboratory, formerly Beth Israel Deaconess Medical Center/Harvard Medical School and currently at the Harrington Discovery Institute, Cleveland.

All mouse care, handling, and experimental procedures were approved by the Mayo Clinic Institutional Animal Care and Use Committee in accordance with National Institutes of Health guidelines. Mice were housed in standard Plexiglass cages with rodent chow and water available *ad libitum*, unless otherwise stated. The colony room was maintained at a constant temperature 24 ± 1°C and humidity 60 ± 2% on a 12-hour light/dark cycle with lights on at 06:00. E27X and R1147C were bred in heterozygous x WT crosses; R579C were bred in heterozygous in crosses. Pups were genotyped at weaning with a unique ear tag and tail biopsy.

PCR primers used were as follows:

E27X F: 5’ / R: 5’ 5’ ATGGAGATCCAGATGGACGA / R: 5’ CCTTCTCGCGCTCGTATTG R579C F: 5’ AGTGGGTGACTTGTAGTGAATG / R: 5’ CGAGTGTTGGCATGGTTTG R1147C F: 5’ TGCCATCTAGGGTTGGAAAC / R: 5’ GAGGCTCATCTTTAGCTCCATAA

### Behavioral phenotyping

Mice were acclimated to the testing room for 30 minutes prior to each assessment. For open field test, elevated plus maze, and rotarod experiments, approximately 10 males and females of heterozygous and homozygous (R579C) patient variant strains were tested at 3- and 6-months of age. Gait analysis was collected at a single timepoint in mice at 3-months of age, while treadmill performance was assessed at 12-months of age. Experimenters were blinded to genotype, and experiments were performed by in-crossing heterozygous or heterozygous and FVB wild-type animals.

### Elevated Plus Maze

Animals were evaluated for anxiety-like behaviors using an elevated plus maze (EPM), a T-shaped maze raised 74 cm above the ground, consisting of two open (35 cm x 6 cm) and two closed (35 cm x 6 cm x 22 cm walls) arms connected by a central zone platform (6 cm x 6 cm) (Med Associates) (33). Animals were placed in the center of the platform and allowed to explore over a 10-minute period. Movement and location were tracked overhead by nose, center, and tail points using Ethovision-XT (Noldus). After each session, mazes were sanitized to remove olfactory cues and organic waste.

### Treadmill assay

The treadmill assay was performed to measure exercise capacity and endurance (34,35). Mice were trained on the treadmill for 3 sessions, performed once per day, prior to data collection. Each mouse was placed on the treadmill surface at initial speed of 5 meters/minute (m/min) with speed increased by 2 m/min every 2 minutes until a maximum speed of 9 m/min was reached (total of 6 minutes per training session). To encourage continued walking, an electrified metal grid, located at the rear of the moving belt, delivered a mild aversive stimulus when mice paused in walking. On the test day, mice were placed on the treadmill and ran at increasing rates (starting at 5 m/min and again increasing by 2 m/min every 2 minutes) until exhaustion.

Exhaustion was defined by the mouse remaining on the electric grid for 2-5 seconds, at which time the electric pulse was turned off to allow the mouse time for rest and recovery. Results were recorded as total time running before reaching exhaustion (36).

### Open Field Test

To evaluate locomotor activity and anxiety-like behavior, the amount of time spent in seconds (s) and distance traveled (cm) in the center and peripheral zones were compared by genotype, age, and sex using the open field test. For all open-field test (OFT) experiments, ENV-510 test environment equipped with infrared beams and Activity Monitor (Med Associates) evaluated spontaneous motor activity during the open field test. Mice were placed in a Plexiglas box (27 x 27 x 20.3 cm) and allowed to explore the chamber for 10 minutes (37).

The data were recorded by each beam break as one unit of exploratory activity using the activity monitoring software (Med Associates).

### Rotarod assay

The rotarod assay tested motor coordination and balance by genotype, sex, and age (3- and 6-months old). Mice were placed on the rotating cylinder with an initial speed of 4 revolutions per minute (rpm). Over an acceleration time of 5 minutes, the rod reached a maximum speed of 40 rpm, where it remained for the duration of the trial. Trials ended when the mouse fell off the rod, performed a passive rotation, or after reaching the cutoff time of 5 minutes. Each mouse was tested 3 times, and mean latency to fall was recorded and averaged (38).

### Gait analysis

Analogue analysis was performed to quantify gait mechanics. Footprints were recorded by coating the bottom of the mouse’s feet with non-toxic washable paint (one color for fore paws and a contrasting color for the hind paws) before allowing the mouse to walk across a strip of paper cut to 3.5 in x 15 in placed underneath a clear acrylic tunnel (39). Mice were encouraged to walk through the tunnel by placing enrichment items from the housing cage at the opposite end as incentive. Results were reported as average stride length (distance between center of foot pad between steps for each forefoot, left and right) and average stride width (distance between center of foot pad of left and right forefoot).

### Hindlimb clasping

The hindlimb clasping assay was conducted as previously described (40). Briefly, 3-month-old male and female mice were gently held in place by the tip of their tail and suspended approximately 40 cm above a tabletop. Movements were recorded for 15 seconds, ensuring that both hindlimbs and the abdomen were clearly visible to the camera, throughout the recording period. Clasping events were recorded for one or both hindlimbs. Any clasping behavior lasting 3 or more seconds was scored as 1 (“clasping”), or 0 (“no clasping”). The experimenter holding the mice and scoring the videos was blinded to genotype, which was determined after the completion of the assay.

### Brain weight measurements

Mouse brains were isolated following transcranial perfusion. Each brain was weighed using a calibrated analytical scale, and total brain weight (mg) was recorded.

### Statistical analysis of behavioral data

Normality data was tested using the Shapiro-Wilk test and significant differences between groups were evaluated using one-way ANOVA or Kruskal Wallis test (parametric), multiple comparisons using Tukey’s post hoc test for multiple comparisons, assuming an interval of 95% confidence (p <0.05). The influence of age and sex were investigated, and analyses were performed with GraphPad Prism software.

### Protein sample preparation

Immediately following standard CO_2_-mediated euthanasia, whole brain was flash frozen and stored at −80°C for at least 24-hours before being resuspended in ice-cold lysis buffer (50 mM Tris-HCl pH 7.4, 150 mM NaCl, 1 mM EDTA, 1% Triton-X 100, 1:100 Halt Protease and Phosphatase Inhibitor Cocktail), ground utilizing a T25 digital ULTRA-TURRAX disperser (IKA), and then centrifuged for 10 minutes at 17,000g and 4°C. The supernatant was retrieved, and total protein concentration level was measured using the Smith assay (Thermo Fisher Scientific A55864).

### Quantification of JIP3 levels in 3-month-old mouse brains by western blot

Equal amounts of protein, 20 µg, were separated by 4-15% SDS-PAGE (BioRad 5671083 & 5671084) and transferred to PVDF membranes (BioRad 1620177). The membranes were blocked in 5% nonfat milk for 1 hour and then incubated at 4°C overnight with the following antibodies: rabbit anti-JIP3 targeting the 171-248 amino acid region of the JIP3 protein (1:1000 dilution, Proteintech 25212-1-AP) and rabbit anti-GAPDH (1:1000 dilution, Cell Signaling 5174S). Membranes were washed with TBST (Tris buffered saline, 1% Tween-20) buffer, incubated with HRP-conjugated secondary antibody (Santa Cruz sc-2357) at room temperature for 1 hour, and then briefly incubated, 3-5 minutes, with a chemiluminescent substrate (Thermo Fisher Scientific 34577) for band visualization. An iBright Imaging System (Invitrogen) was used for detecting and quantifying band intensity using densiometry analysis with the iBright Analysis Software (Thermo Fisher Scientific). Ratios were calculated by dividing JIP3 intensity GAPDH intensity, normalized to the average wild-type ratio. Data was checked for normality and a Kruskal Wallis ANOVA with Dunn’s multiple comparison test to determine significant differences compared to wild-type controls.

## RESULTS

### Phenotype-genotype relationships in clinically assessed NEDBA patients

All individuals across all three genotypes exhibited delay in gross motor skills and language (**Table 2**, **Figure 1A**). Microcephaly was not observed in the E27X individual but was present in 80% and 50% of the R578C and R1146C individuals, respectively. Delays in fine motor skills were observed in the E27X individual and present in the majority of R578C (80%) and RH1146C (83.3%) individuals. Hypotonia was observed in the person with the E27X variant and most (66.7%) of the R1146C individuals but was uncommon (20%) in R578C individuals. While not a ubiquitous manifestation, hypertonia was a distinctive feature (60%) of the R578C individuals, rarely observed (8.3%) in the R1146C individuals and absent in the E27X person. Among the individuals with the R578C variant, 40% exhibited tremor, myoclonus, and dystonia, all features that were not present in the R1146C cohort. The single E27X person exhibited dystonia and ataxia.

**Figure 1.**
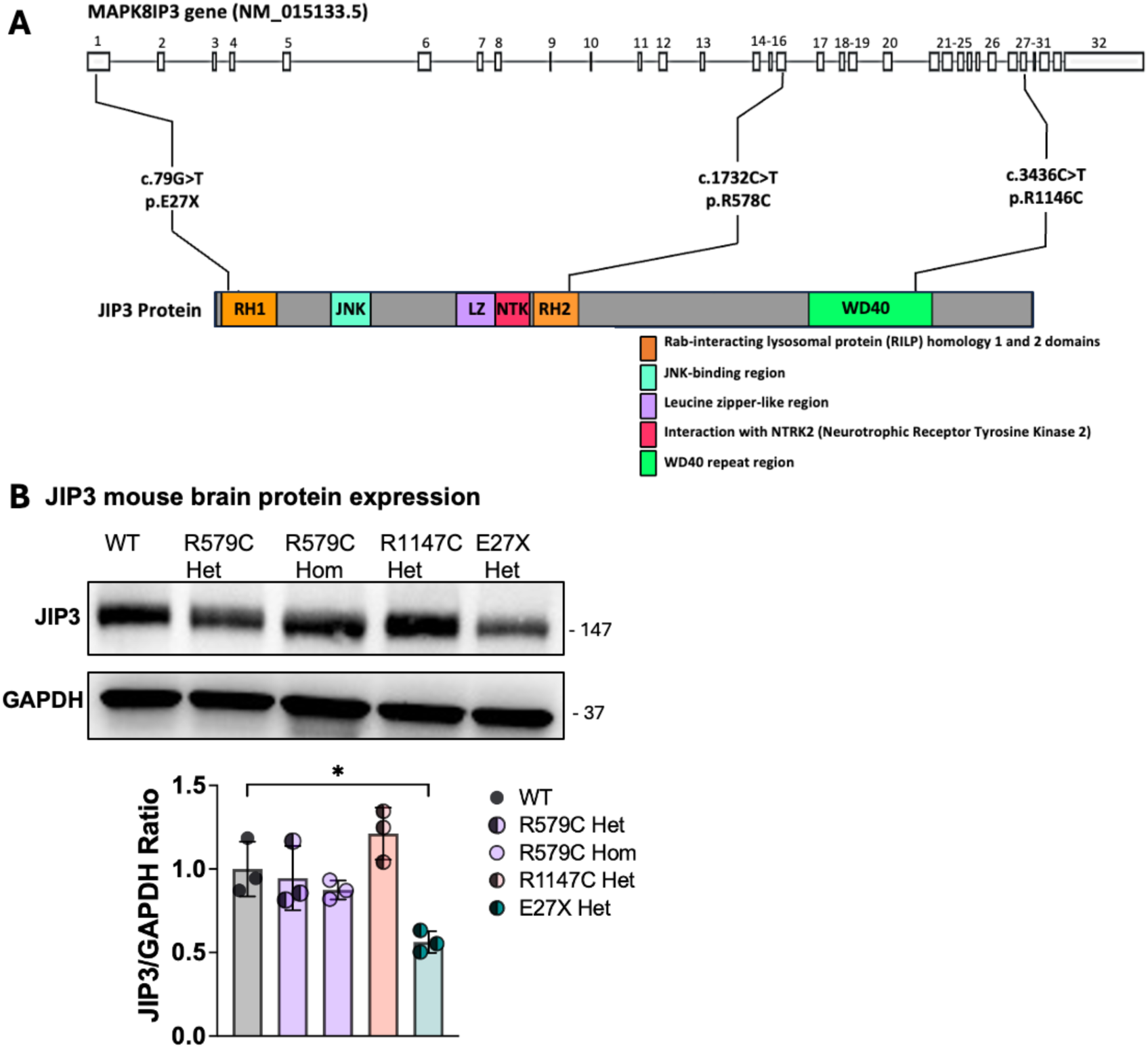
Location, description, and protein expression of *MAPK8IP3* patient variants examined in this study. **(A)** The human *MAPK8IP3* gene (NCBI references number NM_015133.5) has 32 exons. DNA and amino acid changes, provided by an ongoing natural history study, are provided for each pathogenic variant in this study. Variants are located across three individual JIP3 protein domains (colored boxes). These variants represent a subset of all *MAPK8IP3* variants and were selected due to their location in distinct protein domains. Gene information was obtained from Ensembl. JIP3 and GAPDH protein expression from the brains of 3-month old female mice with quantified protein expression provided as a ratio to GAPDH expression (**B**). JIP3 and GAPDH protein levels were quantified using iBight software and statistical analysis was conducted using Kruskal Wallis with Dunn’s test for multiple comparisons.

For the cognitive domains, intellectual disability was common, and observed in the E27X individual, 60% of R578C individuals, and 66.7% of the R1146C individuals. ADHD and ASD were reported in the E27X individual and 50% (ADHD) and 25% (ASD) of the R1146C group but neither were observed in R578C individuals. Anxiety was present in the E27X individual and moderately frequent in the R578C (40%) and R1146C (33.3%) groups. Sensory integration disorder was overall less common, observed only in 33.3% of R1146C group, 20% of R578C group and absent in the E27X individual.

Protein homology of canonical human (transcript ENST00000610761.2) and mouse (transcript ENSMUST00000088345.12) (**Table 1**) protein sequences show 94% identity in a BLASTp alignment and all three patient variants (E27X, R578C, R1146C) are conserved across species (**Supplemental Figure 1**), and nucleotide changes and restriction enzyme sites for genotyping are shown (**Supplemental Figure 2**).

**Table 1.**
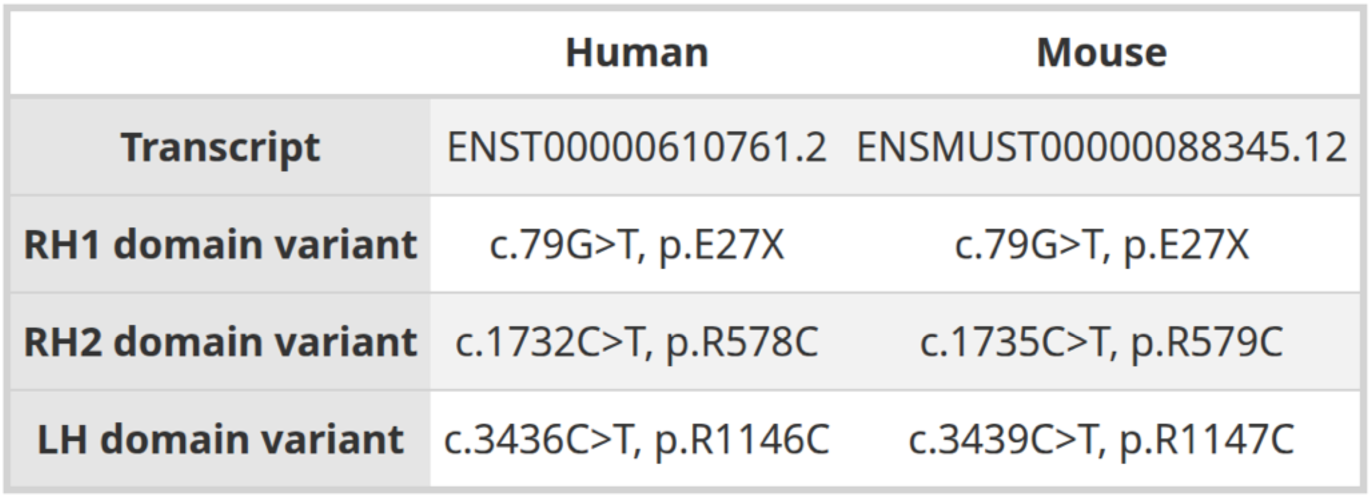
Human and mouse *MAPK8IP3* transcripts, variant information and variant domain location. . Transcript information is provided by Ensembl. Corresponding variant location in mouse ortholog for knock-in models is provided.

**Table 2.** Genotype-associated clinical features in *MAPK8IP3*-related NEDBA individuals. Percentages of individuals exhibiting neurological symptoms within variant groups are provided for each genotype. Acronyms: Attention-deficit hyperactive disorder (ADHD).

|  |  | Variant NM_015133.5 |  |  |
| --- | --- | --- | --- | --- |
|  |  | p.E27X | p.R578C | p.R1146C |
|  |  | *n* = 1 | *n* = 5 | *n* = 12 |
|  | Microcephaly | 0.0% | 80.0% | 50.0% |
| Cognitive Domains | Intellectual disability | 100.0% | 60.0% | 66.7% |
|  | Delay in language | 100.0% | 100.0% | 100.0% |
|  | Autism | 100.0% | 0.0% | 25.0% |
|  | Sensory integration disorder | 0.0% | 20.0% | 33.3% |
|  | ADHD | 100.0% | 0.0% | 50.0% |
|  | Anxiety | 100.0% | 40.0% | 33.3% |
| Motor Domains | Delay in gross motor | 100.0% | 100.0% | 100.0% |
|  | Delay in fine motor | 100.0% | 80.0% | 83.3% |
|  | Hypotonia | 100.0% | 20.0% | 66.7% |
|  | Hypertonia | 0.0% | 60.0% | 8.3% |
|  | Dystonia | 100.0% | 40.0% | 0.0% |
|  | Myoclonus | 0.0% | 40.0% | 0.0% |
|  | Tremor | 0.0% | 40.0% | 0.0% |
|  | Ataxia | 100.0% | 0.0% | 0.0% |

### Mouse Embryonic lethality

E27X and R1147C homozygous mice were embryonic lethal, and no homozygous pups were viable at birth.

### Decreased Jip3 protein levels in E27X mice

To measure the impact of missense or truncating variants in *Mapk8ip3* on protein expression in the brain, we measured the levels of JIP3 in whole brain lysates of 3-month-old knock-in female mouse brains. As expected, the brains of mice heterozygous for E27X truncating variant showed approximately 50% lower JIP3 protein levels compared to wild type JIP3 levels were comparable to wild type in heterozygous and homozygous in R579C brains and slightly elevated, although not significantly, in heterozygous R1147C brains (**Figure 1B**).

### Mouse Behavioral deficits

The open field test (**Figure 2A-C and 3C**) and elevated plus maze (**Figure 3A-B** and **Supplemental Figure 3**) were used to examine motor activity and anxiety-like behavior across all mouse genotypes, sexes, and age ranges (3- and 6-months old). Significant differences and trends in locomotor activity and behavior across these two assays were largely consistent. Only stride width differences were observed in heterozygous and homozygous R579C and heterozygous R1147C for gait analysis (**Figure 4C** and **Supplemental Figure 4**).

**Figure 2.**
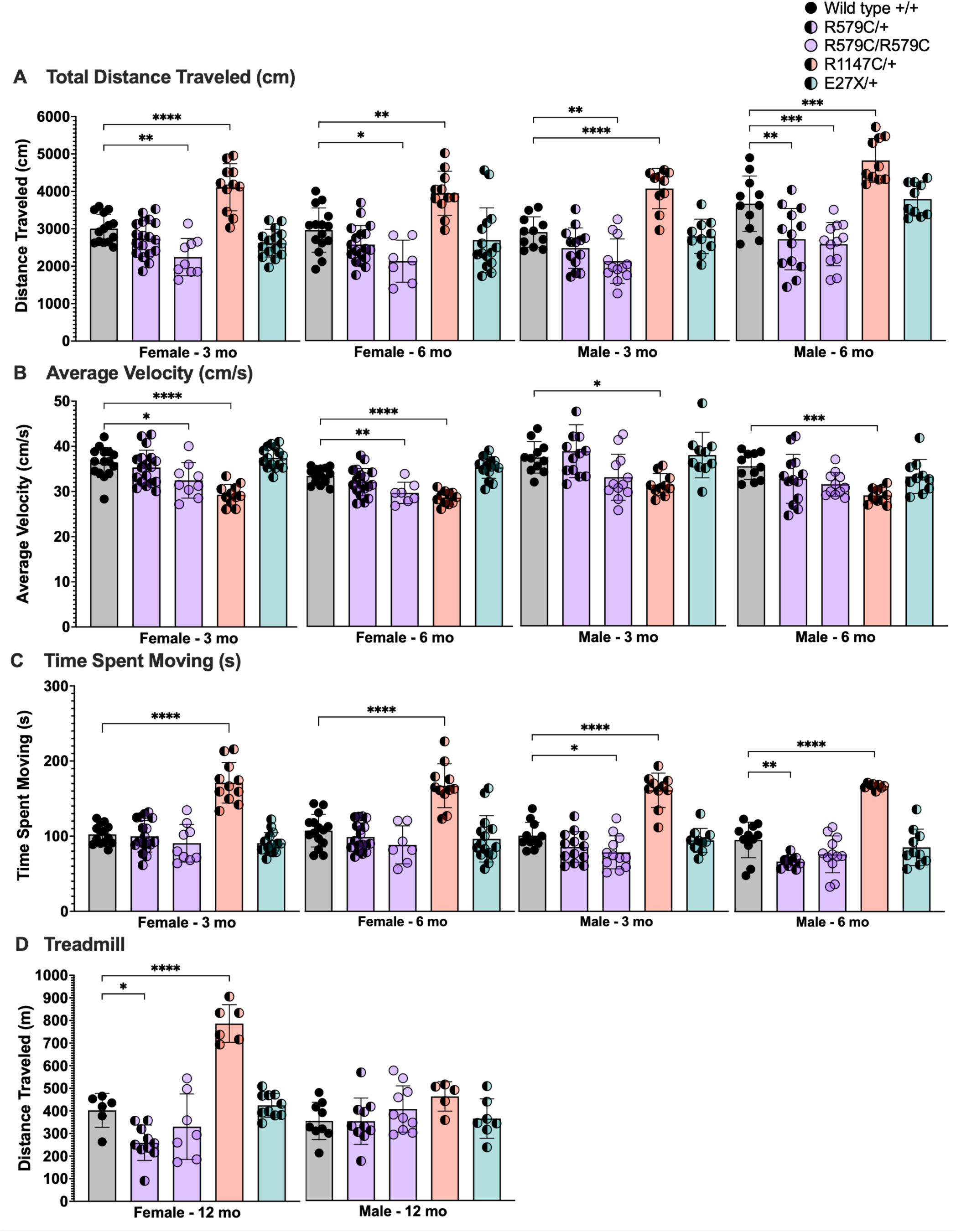
Locomotor phenotypes assessed using the open field test and treadmill assay. **(A)** 3- and 6-month-old female and male homozygous R579C mice and 6-month old male heterozygous R579C mice traveled significantly less distance (cm), whereas 3- and 6-month old heterozygous female and male R1146C mice traveled greater distance (cm), compared to wild-type during the open field test. **B**) 3- and 6-month old heterozygous female R579C mice traveled at decreased average velocities (cm/s), whereas 3- and 6-month old female and male heterozygous R1146C mice traveled at increased average velocities (cm/s), compared to wild-type siblings during the open field test. (**C**) 3-month old homozygous and 6-month old heterozygous male R579C mice spent less time (s) moving, whereas 3- and 6-month old female and male heterozygous R1147C mice spent more time (s) moving compared to wild-type littermates during the open field test. (**D**) 12-month old heterozygous female R579C mice traveled significantly less distance, whereas 12-month old heterozygous R1147C mice traveled significantly greater distances, compared to wild-type siblings during the treadmill assay. Statistical significance was determined using a 1-way ANOVA with Sidak’s test for multiple comparisons, p<0.05=*, p<0.01=**, p<0.001=***, p<0.0001=****.

**Figure 3.**
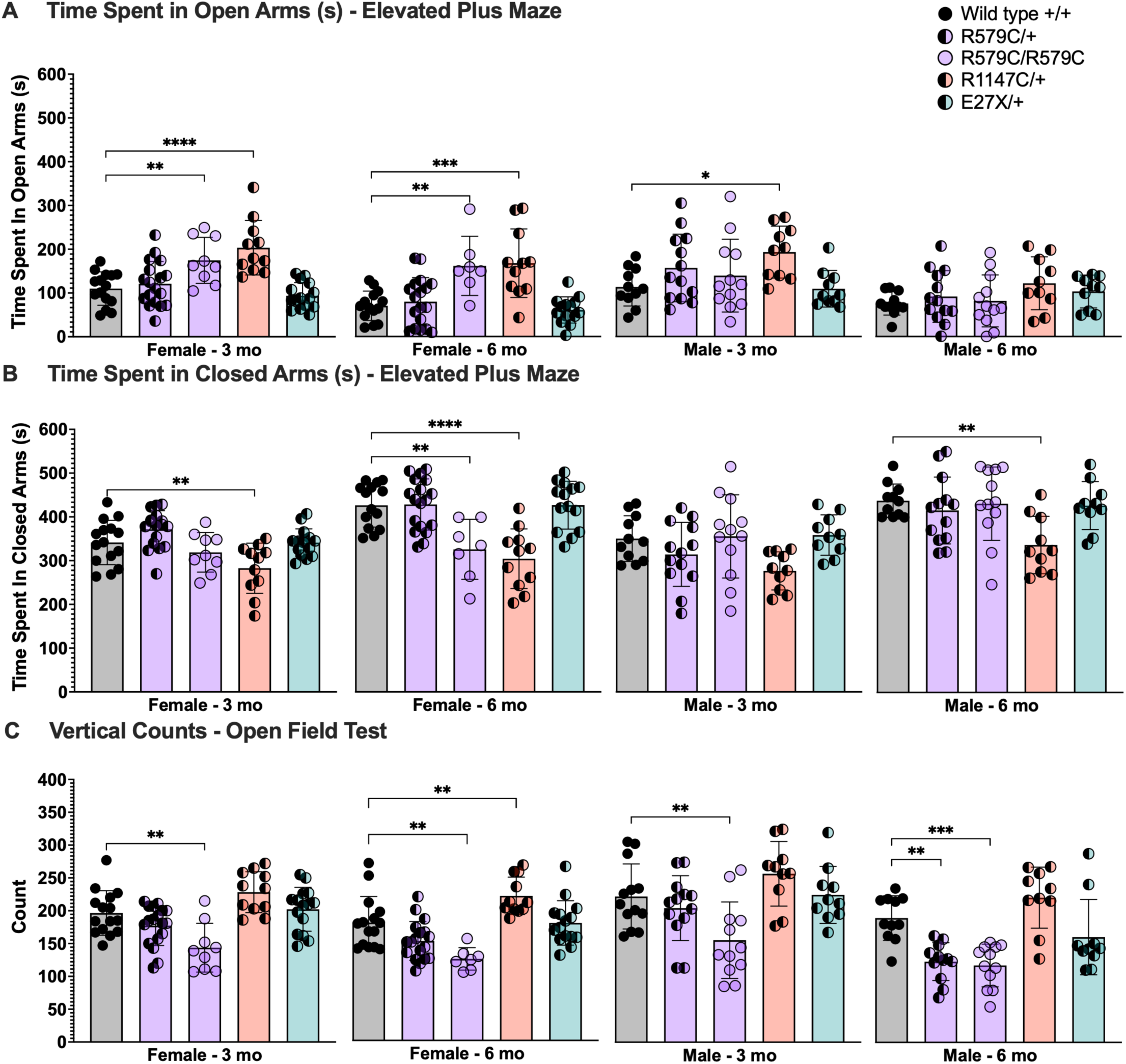
Anxiety-like and exploratory behavior phenotypes. **(A)** 3- and 6-month old homozygous R579C mice and 3- and 6-month old female and 3-month old male heterozygous R1147C mice spent significantly more time (s) in the open arms of the elevated plus maze, compared to wild-type littermates, indicating a decreased anxiety-like phenotype. (**B**) 6-month old homozygous female R579C and 3- and 6-month old female and 6-month old male heterozygous R1147C mice spent significantly less time (s) in the closed arms of the elevated plus maze compared to wild-type siblings. (**C**) 3- and 6-month old female and male homozygous and 6-month old male heterozygous R579C mice displayed less vertical counts, compared to wild-type siblings, indicating less exploratory behavior. In contrast, 6-month old female heterozygous R1147C mice exhibited increased vertical counts, compared to wild-type littermates, indicating increased exploratory behavior. Statistical significance was determined using a 1-way ANOVA with Sidak’s test for multiple comparisons, p<0.05=*, p<0.01=**, p<0.001=***, p<0.0001=****.

**Figure 4.**
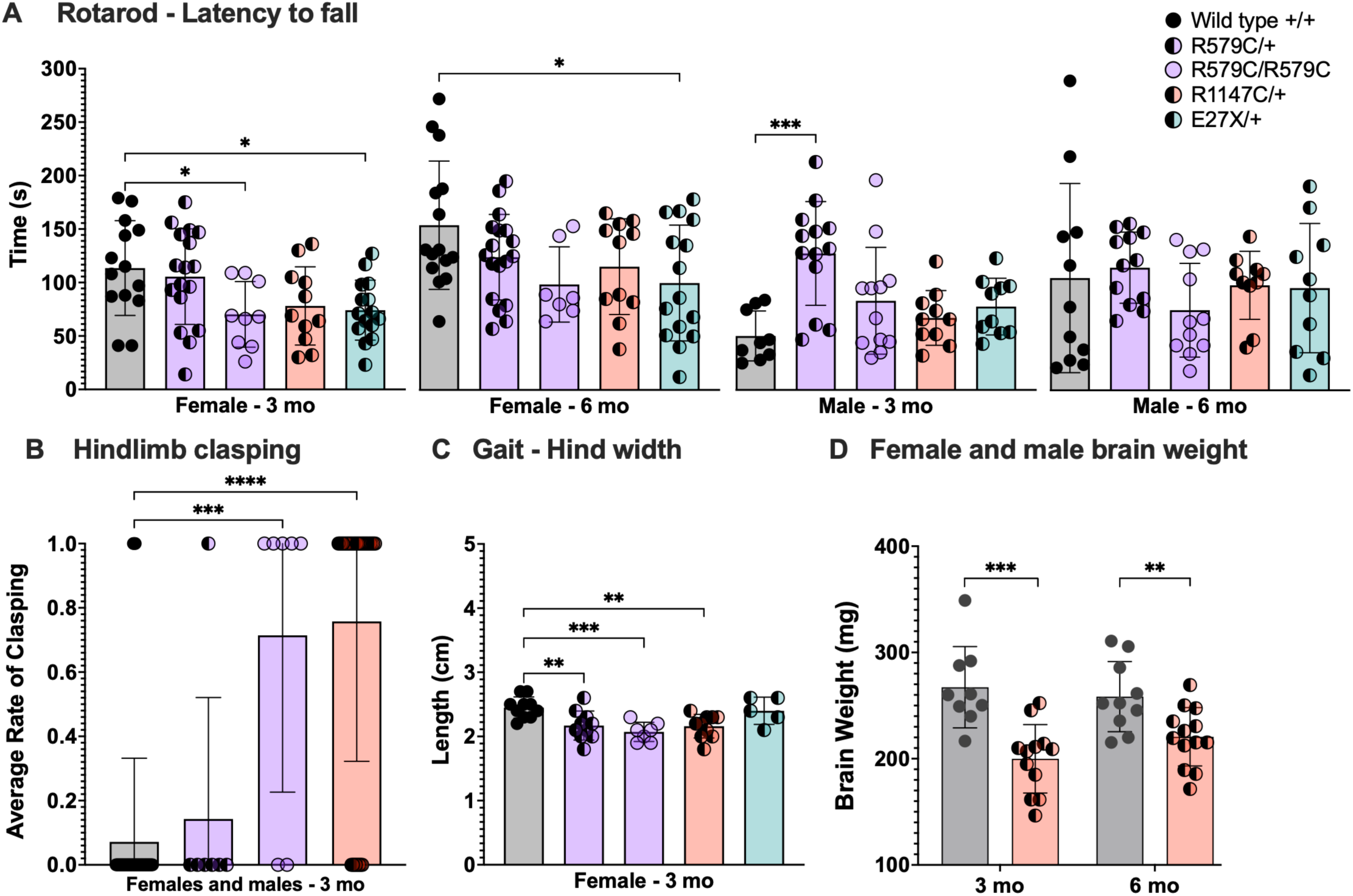
Gross motor and brain weight phenotypes. (**A**) 3-month-old female homozygous R579C mice and 3- and 6-month-old female heterozygous E27X mice exhibited significantly decreased gross motor function in rotarod tests compared to wild-type littermates. 3-month-old heterozygous R579C mice exhibited increased time spent on the rotarod compared to wild-type siblings, however, wild-type sibling male mice exhibited unusually shorter amount of time spent on the rotarod. (**B**) 3-month old homozygous R579C and heterozygous R1146C female and male mice demonstrated significant hindlimb clasping behaviors indicative of impaired gross motor function, whereas wild-type siblings did not. (**C**) 3-month old female heterozygous R579C and R1146C and homozygous R579C mice exhibited increased hind width (cm) in walking gait, no other differences in gate were observed across genotypes (**Supplemental** Figure 3). (**C**) 3- and 6-month old male heterozygous R1146C mice showed significantly reduced brain weight compared to wild-type littermates. Statistical significance was determined using a 1-way ANOVA with Sidak’s test for multiple comparisons, and Mann-Whitney unpaired t-test for two group comparisons p<0.05=*, p<0.01=**, p<0.001=***, p<0.0001=****.

### Decreased locomotor activity in R579C mice

Homozygous R579C male and female mice at 3-and 6-months of age traveled shorter distances (cm) than their wild-type littermates. Heterozygous R579C male mice traveled shorter distances at 6-month of age (**Figure 2A**). Female homozygous R579C mice traveled at slower velocities (cm/s) at both 3-and 6-months of age, compared to wild-type siblings (**Figure 2B**). Male homozygous R579C mice spent less time (s) moving, compared to wild-type siblings, with significant differences observed at 3-months of age (**Figure 2C**). At 12 months, heterozygous R579C female mice traveled at shorter mean distance on the treadmill (**Figure 2D**).

### Increased locomotor activity in R1147C mice

Heterozygous R1147C female and male mice at 3-and 6-months old traveled significantly greater distances (cm), and spent significantly more time moving, at significantly slower velocities, compared to wild-type siblings (**Figure 2A-C**). Twelve-month old heterozygous female mice traveled significantly further distances on the treadmill assay, compared to wild-type siblings (**Figure 2D**).

### Reduced anxiety-like behavior and decreased exploratory behavior in R579C mice

Female 3-and 6-month-old R579C mice spent significantly more time in the open arms of the elevated plus maze (**Figure 3A**). In contrast, R579C female and male homozygous mice at 3- and 6-months old displayed significantly decreased vertical counts, demonstrating less exploratory behavior (**Figure 3C**).

### Decreased anxiety-like behavior in R1147C mice

Heterozygous R1147C female at 3- and 6-months of age and male mice at 3-months old spent significantly greater amount of time in the open arms and less time in the closed arms of the elevated plus maze (**Figure 3A-B**). Female and male 3- and 6-month-old mice on average displayed greater vertical counts, compared to wild-type siblings, with significant differences in 6-month-old female mice (**Figure 3C**).

### Decreased motor skills in female E27X, R579C, and R1147C mice

Heterozygous E27X female mice at 3- and 6-months old displayed significantly decreased latency to fall as measured by time in seconds (s) spent on the rotarod assay, an assay for gross motor coordination (**Figure 4A**). No significant or trending distinctions were observed in heterozygous E27X male mice, compared to wild-type siblings. Homozygous R579C female mice at 3- and 6-months old displayed decreased average time before falling, with significant differences observed in 3-month-old females. Of note, 3-month-old wild-type male mice showed earlier latency to fall than the rest of the wild-type groups, suggesting underperformance in the control group. Latency to fall average values in 3-month-old homozygous R579C males, as well as heterozygous R579C, R1146C, and E27X males were comparable to 3-month-old females of the same genotype. Heterozygous R1147C females at both 3- and 6-months old on average showed a slight decrease in time spent on the rotarod but was not significantly different from wild-type siblings (**Figure 4A**). Hindlimb clasping is another assay that measures motor dysfunction. R579C homozygous and R1147C heterozygous combined male and female mice at 3-months old exhibited significant increased average rates of hindlimb clasping. Heterozygous R579C mice exhibited normal low rates of hindlimb clasping, similar to wild-type controls (**Figure 4B**). Data were not collected for heterozygous E27X mice.

### Decreased brain weight

Heterozygous male and female R1146C mice at 3- and 6-months of age exhibited decreased brain weight at time of perfusion (**Figure 4D**).

## DISCUSSION

This study integrates neurological phenotypes of individuals with pathogenic *MAPK8IP3/*JIP3 variants with longitudinal motor and behavioral analyses in corresponding murine models across developmental stages, providing insight into genotype-phenotype relationships and domain-specific functions of JIP3. By comparing a truncating variant (E27X) with missense variants (R578C and R1146C) located in distinct JIP3 functional domains, we identify both shared and unique neurodevelopmental impairments and motor and behavioral phenotypes.

### Shared neurological phenotypes across genotypes

Global language and gross motor delays were observed across all genotypes in people, suggesting that JIP3 serves fundamental roles early in neurodevelopment across functional brain regions. Because early language and motor milestones depend on axonal outgrowth, synapse formation, and long-range connectivity, JIP3 dysfunction may alter neurodevelopment, producing a common early delay that later gives rise to domain- and dosage-dependent motor and behavioral phenotypes. This is supported by studies in which JIP3 knockout mice were embryonic lethal and showed telencephalic anomalies (2).

Consistent with delays in gross and fine motor development in humans, reduced performance on the rotarod assay was seen in homozygous R579C and heterozygous E27X mice. Likewise, homozygous R579C and heterozygous R1147C mice exhibited prominent hindlimb clasping. These findings suggest deficits in descending motor drive and network-level coordination, potentially involving cortico-cerebellar and basal ganglia-thalamocortical regions, rather than primary impairment of basic locomotor pattern generation.

However, studies examining circuit-level differences are needed to support this hypothesis. Despite the motor abnormalities, only stride width differences in gait were observed in homozygous and heterozygous R579C and heterozygous R1147C mice.

Both R579C and R1147C mouse models exhibited reduced anxiety-like behavior, as reflected by increased time spent in the open arms of the elevated plus maze. Although this phenotype should not be interpreted as a direct measure of affective state, it provides evidence for altered behavioral inhibition and risk assessment governed by distinct brain regions including prefrontal, limbic, basal ganglia, and brainstem networks, secondary to primary motor and coordination impairment. Together, these shared findings suggest higher-order motor and behavioral circuit dysfunction across JIP3 variants.

### JIP3 protein domain-specific neurological features

In humans, the truncating E27X variant (albeit, in the single individual examined thus far), is associated with a broad neurodevelopmental phenotype including hypotonia, fine motor delay, ataxia, dystonia, and prominent neurobehavioral features such as intellectual disability, ADHD, ASD, and anxiety. Consistent with the predicted severity of a truncating allele, homozygous E27X mice were embryonic lethal. This underscores that JIP3 function is critical during early development and establishes a clear gene-dosage threshold for viability. In contrast, heterozygous E27X females exhibited reduced rotarod performance without detectable gait abnormalities, indicating preserved basic locomotor pattern generation but impaired motor coordination, balance, or motor learning (41). Interestingly, the data aligns with the patient data with the E27X person exhibiting ataxia while the R578C and R1146C groups did not. This selective impairment of rotarod performance despite preserved gait suggests that partial loss of JIP3 function preferentially disrupts higher-order motor circuit integration potentially involving cortical, cerebellar, and basal ganglia networks (41,42).

Human variant groups exhibited distinct abnormalities in muscle tone and movement. Hypotonia was present in the E27X individual and the majority of the R1146C group but was uncommon in the R578C group. Hypotonia is consistent with reduced descending motor drive, potentially reflecting impaired corticospinal or cortico-cerebellar output, delayed maturation of long-range motor pathways, or diminished activation of spinal motor neurons (43). In contrast, hypertonia was a defining feature of the R578C cohort but rarely observed in R1146C individuals. Tremor, myoclonus and dystonia were likewise present in the R578C cohort but absent in those with the R1146C variant.

The combination of hypertonia, tremor, and dystonia in the R578C group suggests disinhibition within motor control circuits, particularly those involving basal ganglia and brainstem motor pathways, consistent with established models of dystonic physiology (43,44). Homozygous R579C female mice exhibited decreased locomotor activity, impaired rotarod performance, and prominent hindlimb clasping across development. These features suggest network-level dysfunction rather than primary weakness and may reflect altered development or maintenance of long-range motor projections with high axonal transport demands. However, additional neuropathological examination is needed to support this hypothesis. The basal ganglia, for instance, serves critical roles in regulating muscle tone and movement initiation. Given the known roles of JIP3 in axonal transport, functional disruption of the long, highly branched axons of the nigrostriatal pathway or other central motor circuits is biologically plausible (45). Whether these abnormalities arise from impaired circuit formation during development or from reduced stability and maintenance of projections over time remains unclear.

Overall, the concordance between human and mouse phenotypes supports the interpretation that disruption of the RH2 domain of JIP3 preferentially alters motor circuit regulation, likely involving supraspinal motor circuits such as basal ganglia or brainstem pathways, rather than simply reducing motor output.

Similarly, the R1147C mouse phenotypes also closely parallel phenotypes in R1146C people, who more frequently exhibited neurobehavioral features such as attentional dysregulation and hyperactivity, with fewer hyperkinetic movement abnormalities. The R1147C mouse model exhibited hyperactive locomotor activity, increased exploratory behaviors, and preserved rotarod performance, indicating intact motor coordination and strength. Despite this relative preservation of coordinated motor output, R1147C mice displayed prominent hindlimb clasping and reduced brain weight, suggesting altered motor circuit development or maturation rather than primary motor weakness.

Preserved motor coordination with hypotonia in humans suggests impaired descending motor drive, potentially suggestive of corticospinal tract development abnormalities, reduced excitatory input onto spinal motor neurons, or spinal motor dysfunction. However, additional studies closely examining the electrophysiology and neuropathophysiological differences are needed to confirm this line of thought and provide important implications for therapy. Together, these findings suggest that disruption of C-terminal WD40 domain of JIP3 preferentially affects the development or maintenance of long-range corticospinal projections or motor neuron connectivity, leading to diminished motor drive and hypotonia rather than disinhibition-driven hypertonic phenotypes. These interpretations are also consistent with the substantial demands for cargo trafficking in corticospinal and spinal motor neurons, whose axons are among the longest in the nervous system. Together, these divergent motor phenotypes across JIP3 variants suggest that disruption of specific protein domains differentially impact aspects of motor control, rather than producing uniform impairments.

### JIP3 dosage effect in mouse models

Homozygous E27X and R1147C mice were embryonic lethal, whereas homozygous R579C mice were viable. This indicates that distinct JIP3 domains contribute differentially to developmental viability. Lower JIP3 protein levels were expected with the E27X truncating variant, resulting in a haploinsufficiency model. Interestingly, homozygous and heterozygous R579C mice showed greater motor and behavioral deficits relative to heterozygous E27X truncating variant, which could be the outcome of a toxic gain of function mechanisms (20). Disruption of the C-terminal WD40/cargo-binding domain (R1147C) appears incompatible with early development, suggesting that cargo recognition, scaffolding, or signaling functions of JIP3 are essential for neuronal survival and circuit formation. In contrast, disruption of the RH2 motor-interaction domain (R579C) permits survival but results in postnatal motor dysfunction, indicating that impaired motor protein–cargo coupling is tolerated during development but compromises circuit regulation post early development. These findings support a domain-specific hierarchy of JIP3 function in which cargo binding is required for viability, whereas motor protein interaction primarily shapes postnatal motor phenotypes. However, additional neuropathological studies are needed to decipher whether these differences are shaped by developmental deficits or reduced neuronal maintenance over time.

This study employs well established behavioral assays of motor function as a first step to introduce novel mouse models of neurodevelopmental disorders with cognitive and motor manifestations. We propose that key clinical features are recapitulated, with hyperactivity and exploratory behavior in R1147C, locomotion deficits in R579C, and ataxia in E27X. Beyond the limitations of the inability of murine system to fully recapitulate the human condition, this study focused predominantly on motor domains at mouse ages approximating young to mid adulthood. An additional limitation of a single heterozygous E27X patient should also be noted. Consistent with the human data, we did not find a degenerative phenotypic pattern in mice up to 12 months. This study provides initial comparison and lays the groundwork for future studies to characterize earlier time points to capture developmental trajectory of theses phenotypes. Furthermore, cognitive testing will provide a more comprehensive view of the extent to which variants impact cognitive function in mice, which is the ubiquitous feature of individuals with NEDBA.

## Conclusions

The convergence of reduced anxiety-like behavior with either hypertonic movement disorders (R579C) or hyperactivity and hypotonia (R1147C) suggests that disruption of JIP3 function may impair inhibitory control at the circuit level, yielding variant-specific motor outcomes with some shared features. However, additional studies examining the neuropathology of individual variants are needed to support this hypothesis. These findings support a domain- and dosage-dependent model of JIP3 function in which distinct protein regions make different contributions to developmental viability and motor circuit regulation. Disruption of the C-terminal WD40/protein-interaction domain phenocopies truncating loss-of-function alleles with respect to embryonic lethality, indicating that JIP3-mediated cargo recognition and scaffolding are essential for early neurodevelopment. In contrast, disruption of the RH2 motor-interaction domain is compatible with survival but leads to postnatal motor dysfunction characterized by impaired coordination and abnormal tone, consistent with altered supraspinal motor circuit regulation. Across variants, preserved gait in mouse models indicates intact basic locomotor pattern generation, while deficits in coordination, tone, and behavior reflect possible higher-order circuit dysfunction. This framework reconciles the range of human phenotypes and establishes JIP3 as a critical regulator of neurodevelopment whose domain-specific functions differentially shape viability, circuit maturation, and motor control.

### Future studies

Given the growing repertoire of JIP3 functions (7,46), future studies should examine how distinct JIP3 variants affect endocytosis, intracellular cargo and organelle transport, growth cone dynamics, and JNK signaling in defined neuronal populations across development. This will help correlate motor phenotypes with subcellular, trafficking defects. In mouse models, targeted approaches could include tract-tracing and diffusion-based analyses of corticospinal and basal ganglia projections, electrophysiological or EMG-based assessments of descending motor drive and motor unit recruitment, and cell-type–specific imaging or proteomic strategies to identify cargo classes most sensitive to WD40-domain disruption. Longitudinal behavioral and circuit-level analyses will be essential for distinguishing primary developmental defects from progressive circuit instability. Together, these approaches will help define variant-specific pathogenic mechanisms, identify common therapeutic targets, and establish these models as platforms for preclinical intervention testing.

## SUPPLEMENTAL FIGURES

**Supplemental Figure 1.**
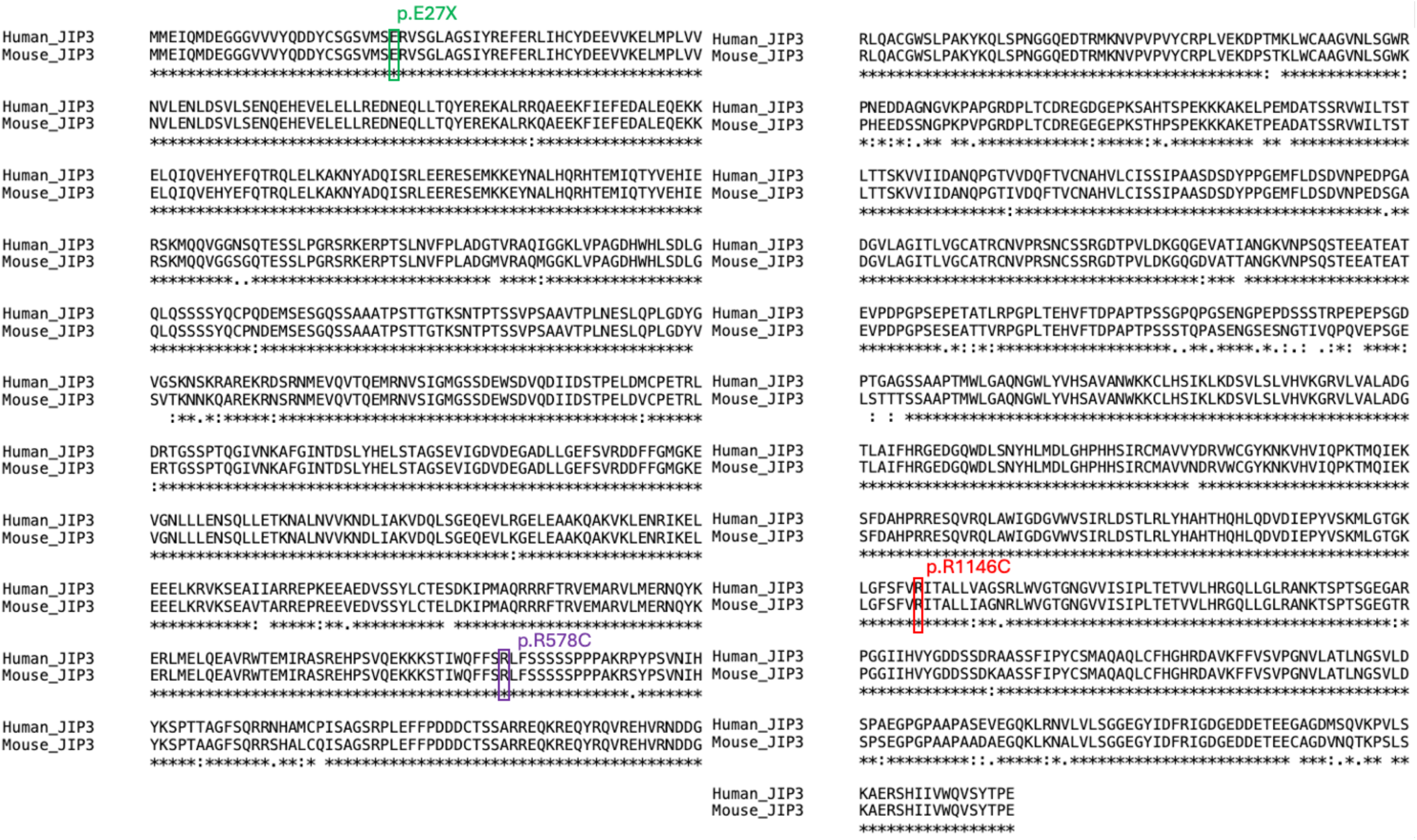
P**r**otein **alignments of JIP3 in human and mouse.** Clustal 2.1 multiple sequence alignment tool confirmed protein homology at variant sites (colored boxes).

**Supplemental Figure 2.**
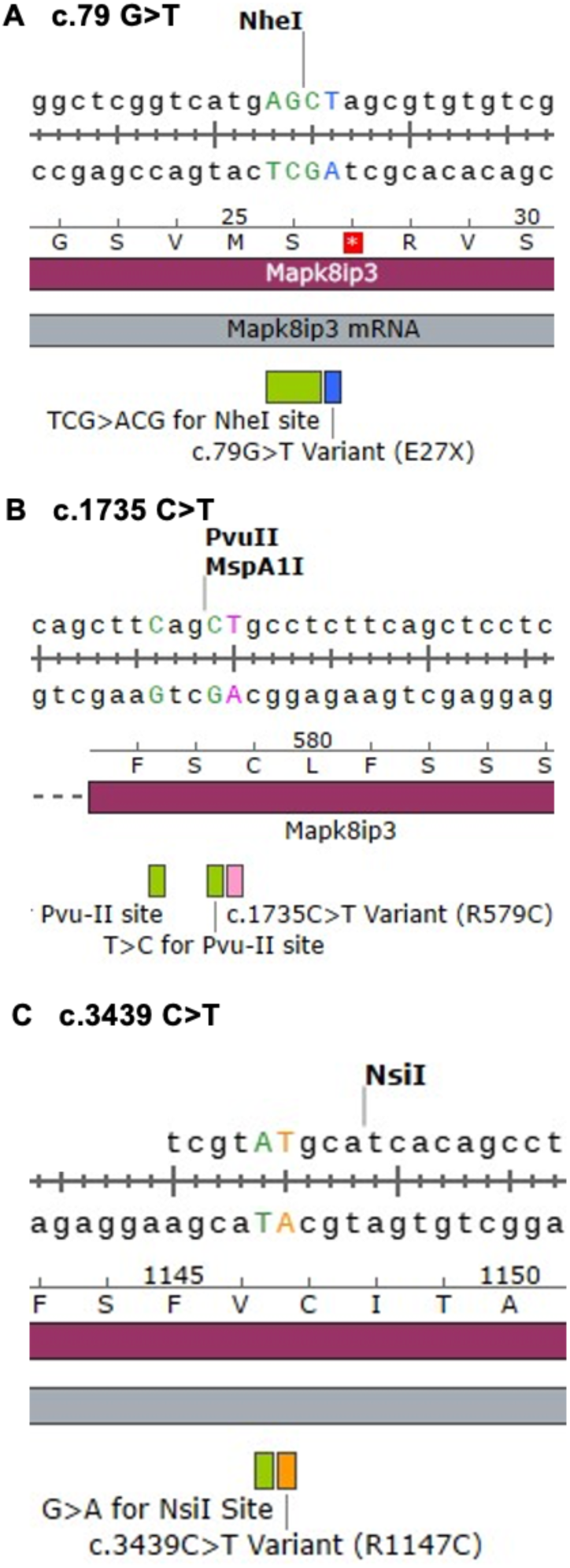
Genotyping of mouse lines shows knock in of patient-variants and restriction enzymes for genotyping.

**Supplemental Figure 3.**
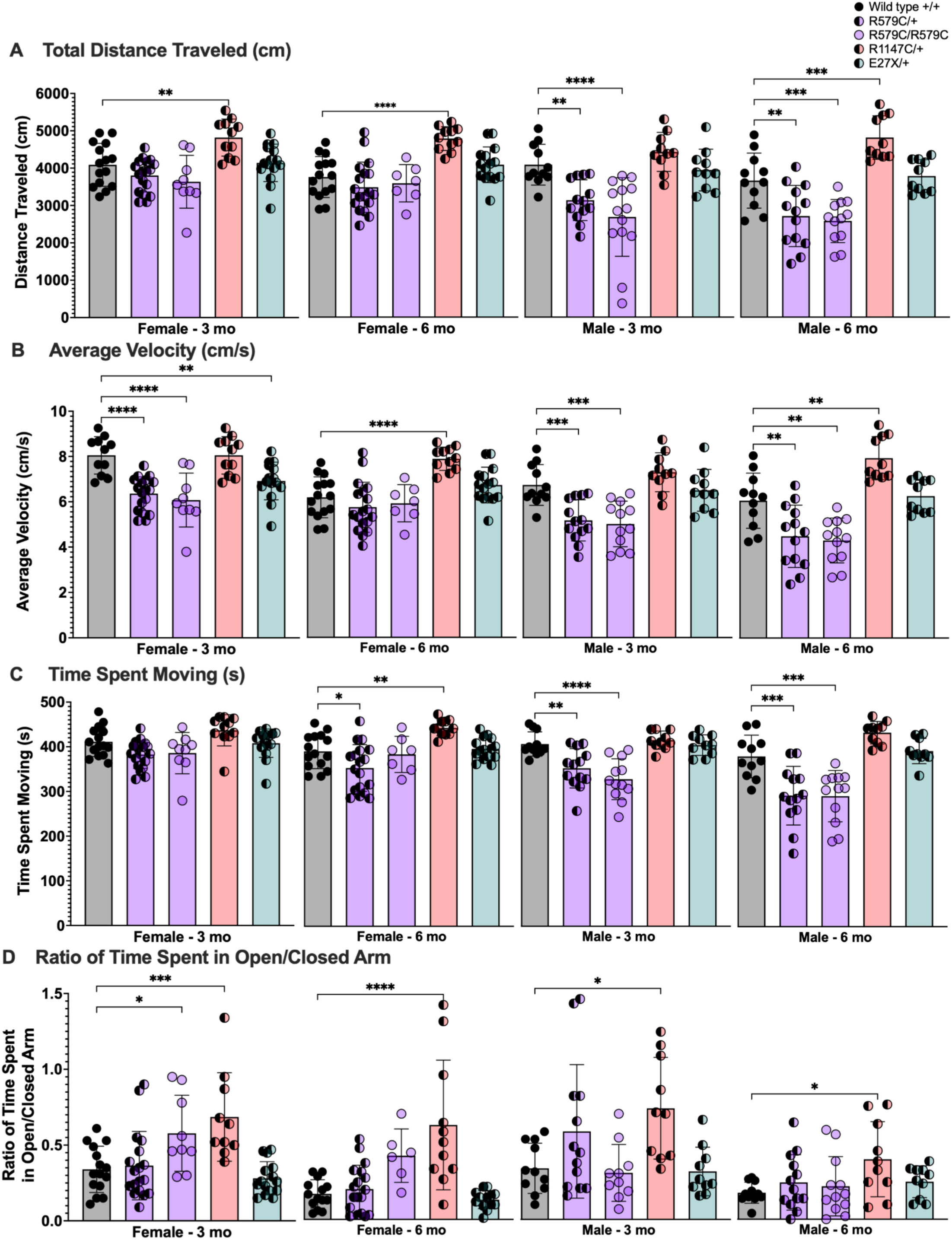
L**o**comotor **phenotypes assessed using the elevated plus maze. (A)** 3- and 6-month old heterozygous and homozygous male R579C mice traveled less distance (cm) compared to wild-type littermates. In contrast, 3-and 6-month old female and 6-month old male heterozygous R1147C mice traveled greater distance (cm) compared to wild-type siblings. (**B**) 3-month old heterozygous and homozygous R579C female, 3- and 6 month-old heterozygous and homozygous R579C male, and 3-month old heterozygous female E27X mice traveled at decreased velocities (cm/s) compared to wild-type siblings. In contrast, 6-month old heterozygous female and male R1147C mice traveled at increased velocities compared to wild-type siblings. (**C**) 6-month old heterozygous female and 3- and 6-month old heterozygous male R579C mice spent less time moving (s) compared to wild-type siblings. 6-month old heterozygous female R1147C mice spent more time moving (s) compared to wild-type siblings. (**D**) 3-month old female R579C and 3- and 6-month old female and male R1147 mice spent a greater ratio of time in the open arm verses the closed arm in the elevated plus maze.

**Supplemental Figure 4.**
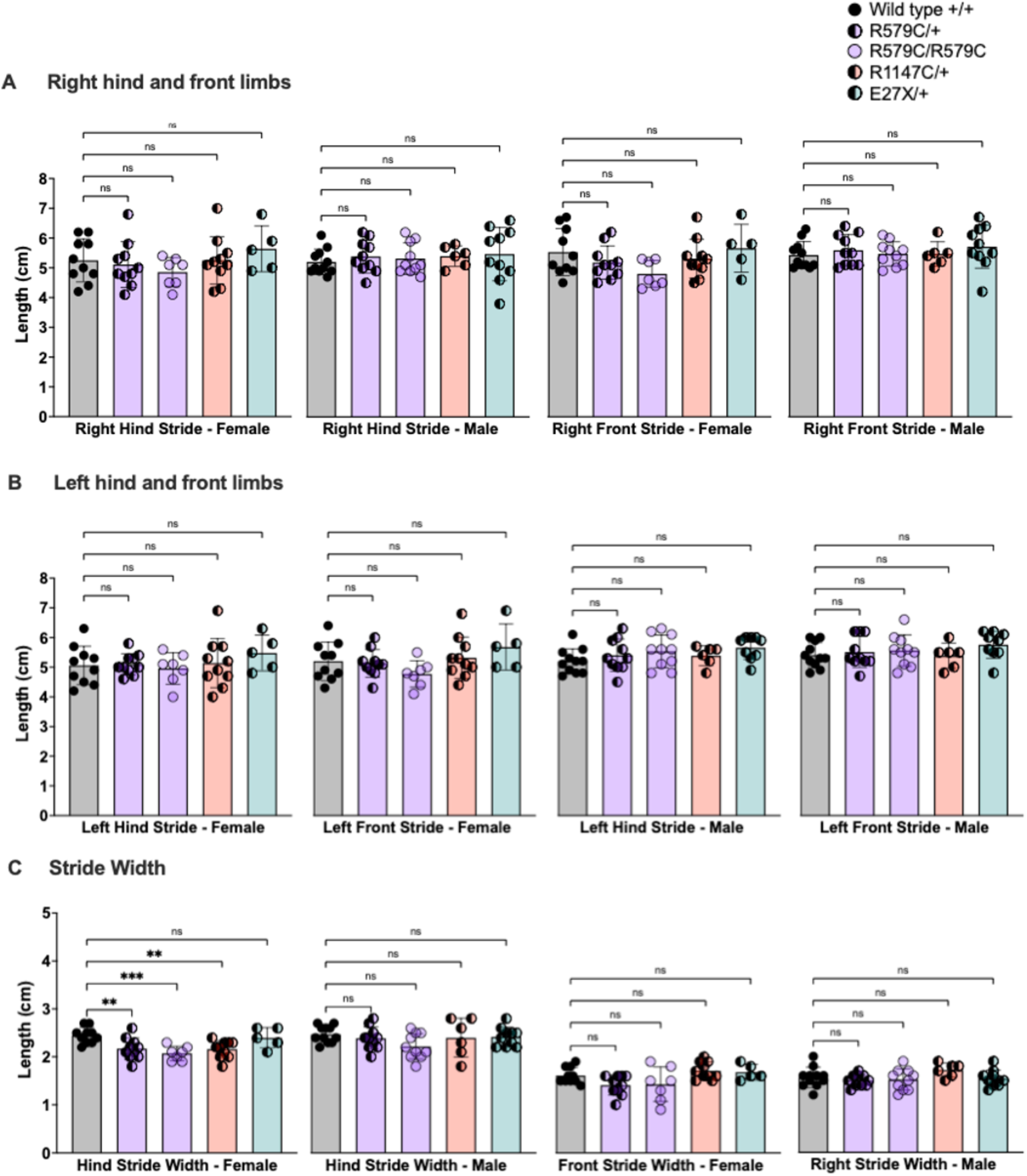
G**a**it **analysis in 3-month old male and female mice. (A)** Right and (**B**) left hind and front limbs stride length (cm) shows no difference, compared to wild-type siblings. **(C)** Female heterozygous and homozygous R579C and heterozygous R1147C show a narrower hind stride width (cm), compared to wild-type siblings. Statistical significance was determined using a 1-way ANOVA with Sidak’s test for multiple comparisons, p<0.05=*, p<0.01=**, p<0.001=***, p<0.0001=****.

